# mTOR inhibits autophagy to facilitate cell swelling and rapid wound repair

**DOI:** 10.1101/2023.10.10.561758

**Authors:** Gordana Scepanovic, Rodrigo Fernandez-Gonzalez

## Abstract

Embryonic wounds repair rapidly, with no inflammation or scarring. Embryonic wound healing is driven by collective cell movements facilitated by the swelling of the cells adjacent to the wound. The mechanistic target of rapamycin complex 1 (mTORC1) is often associated with cell growth. We found that disrupting mTORC1 signalling prevented cell swelling and slowed down wound repair. Catabolic processes, such as autophagy, can inhibit cell growth. Using five-dimensional time-lapse microscopy, as well as pharmacological and genetic manipulations, we demonstrated that the number of autophagosomes decreased during wound repair, suggesting that autophagy must be tightly regulated for rapid wound healing. Quantitative image analysis showed that mTOR inhibition increased autophagy, and that activating autophagy prevented cell swelling and slowed down embryonic wound closure. Finally, reducing autophagy in embryos in which mTORC1 signalling was disrupted rescued rapid wound repair. Together, our results show that mTORC1 activation upon wounding negatively regulates autophagy, allowing cells to increase their volumes to facilitate rapid wound healing.

## INTRODUCTION

The epidermis serves as a barrier that protects an organism from environmental and pathogenic insults. Thus, it is crucial that the epidermis is repaired promptly upon wounding. In both vertebrates and invertebrates, embryos display a remarkable ability to heal wounds rapidly, with no inflammation or scarring (Adzick et al., 1985; Longaker et al., 1990; Whitby and Ferguson, 1991; Martin and Lewis, 1992; McCluskey and Martin, 1995; Martin, 1996; Kiehart et al., 2000; Davidson et al., 2002; Wood et al., 2002). In many species, embryonic wound healing happens at time scales too short for cells to divide. Instead, rapid embryonic wound repair is driven by the polarization of actin and the molecular motor non-muscle myosin II in the cells adjacent to the wound (Brock et al., 1996; Martin, 1996; Kiehart et al., 2000; Davidson et al., 2002; Wood et al., 2002; Hunter et al., 2018). Actomyosin polarization results in the assembly of a contractile cable at the wound edge that coordinates cell movements to draw the wound closed (Wood et al., 2002; Abreu-Blanco et al., 2012; Zulueta-Coarasa, 2018). However, a contractile actomyosin cable is not sufficient for rapid wound repair (Scepanovic et al., 2021b). For embryonic wounds to heal rapidly, the cells adjacent to the wound must swell in a process driven by the local activation of the serine/threonine kinase p38 mitogen-activated protein kinase (Scepanovic et al., 2021b). Cell swelling facilitates contraction of the actomyosin cable by minimizing overall tissue resistance, thus promoting the collective cell movements that drive rapid wound repair.

The mTOR signalling pathway is often associated with cell growth (Liu and Sabatini, 2020). The conserved serine/threonine kinase mTOR assembles a signalling complex, mTORC1, which plays a major role in sensing nutrient availability and controlling cellular homeostasis (Jhanwar-Uniyal et al., 2017). In mice and rats, mTORC1 is required for wound re-epithelialization (Squarize et al., 2010; Huang et al., 2015; Hu et al., 2020), and use of mTOR inhibitors in surgical patients results in wound healing complications (Guilbeau, 2002), suggesting that mTOR signalling plays a key role in wound repair through unknown mechanisms.

Wounding post-embryonic tissues induces autophagy in planarians, *C. elegans, Drosophila*, zebrafish, mice, rats, pigs, and humans (Kakanj et al., 2016; An et al., 2018; Kang et al., 2019; Chavez et al., 2020; Xu et al., 2020; Qiang et al., 2021; Xie et al., 2021; Chen et al., 2022; Deng et al., 2022; Fu et al., 2022; Li et al., 2022). Autophagy is a conserved catabolic “cellular repurposing” system in which intracellular components are packaged into membrane vesicles, autophagosomes, that are delivered to lysosomes, where their contents are degraded (Allen and Baehrecke, 2020). In wounded *Drosophila* larvae, autophagy drives the formation of syncytia, multinucleated cells that facilitate tissue repair (Kakanj et al., 2022). However, exacerbated autophagy can disrupt membrane integrity, and mTORC1 signalling is key to limit autophagy during wound healing. Whether autophagy plays a role in embryonic wound repair has not been investigated.

## RESULTS AND DISCUSSION

### mTORC1 is necessary for rapid wound repair

To determine if mTOR is important for embryonic wound healing, we quantified the dynamics of wound repair in *Drosophila* embryos. We injected embryos expressing the cell-cell junction component E-cadherin tagged with tdTomato (Huang et al., 2009b) and the myosin light chain labeled with GFP (Royou et al., 2004) with 500 µM of rapamycin (Chung et al., 1992), an mTOR inhibitor that primarily targets mTORC1 (Waetzig et al., 2021) (Figure 1A-B, Video S1). We found that wound closure slowed down by 60% in embryos treated with rapamycin with respect to controls (*P* < 0.05, Figure 1C-D). We obtained similar results in embryos expressing an RNAi against the mTORC1 core component *raptor* throughout the embryo (Figure S1A-B): *raptor* RNAi embryos healed wounds 43% slower than *mCherry* RNAi controls (*P* < 0.05, Figure S1C-D). Together, our results indicate that mTORC1 signalling is required for rapid embryonic wound repair.

**Figure 1.**
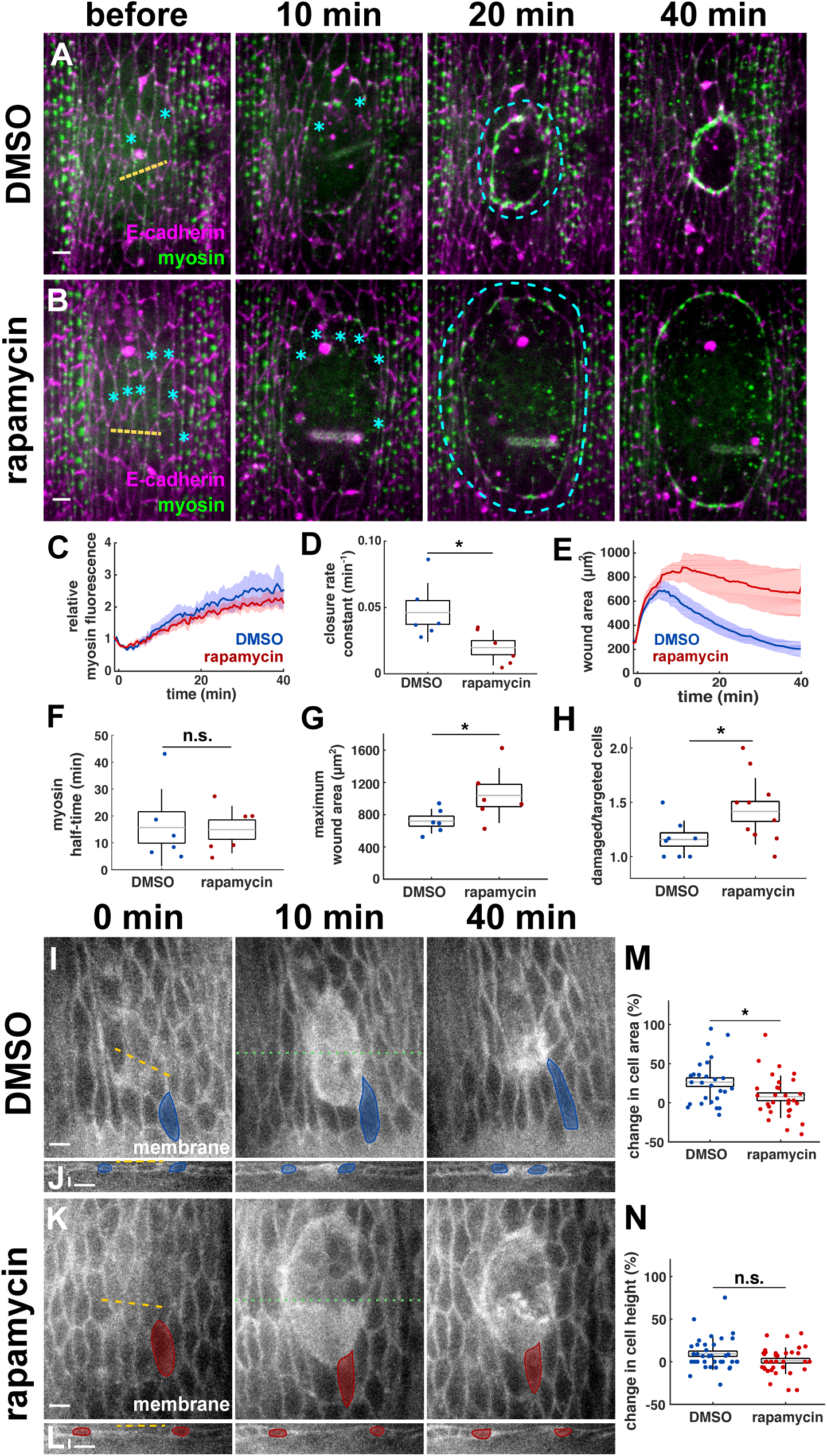
mTOR signalling is necessary for rapid embryonic wound closure. **(A-B)** Wound closure in embryos expressing myosin:GFP (Sqh, green) and E-cadherin:tdTomato (magenta), and injected with DMSO (A), or rapamycin (B). Yellow dashed lines indicate wound sites, cyan dashed lines outline wounds. Cyan asterisks track cells not targeted by the laser but eventually included inside the wound. Time after wounding is shown. Anterior left, dorsal up. Bars, 5 μm. **(C-H)** Wound area over time (C), wound closure rate constant (D), relative mean myosin fluorescence at the wound edge over time (E), half time of myosin accumulation (F), maximum wound area (G), and ratio of damaged cells (inside the myosin cable) to number of cells targeted by the laser (H) for embryos injected with DMSO (blue, *n* = 6 wounds, C-G, or *n* = 8, H) or rapamycin (red, *n* = 6, C-G, or *n* = 10, H). **(I-L)** Apical, XY (I, K), and cross-sectional, XZ (J, L) views of epidermal cells expressing the membrane marker Resille:GFP in embryos injected with DMSO (I-J) or rapamycin (K-L). Sample cells are highlighted in blue (DMSO) and red (rapamycin). Green dotted lines indicate where cross-sections were taken. Time after injection is shown. Anterior, left. Bars, 5 μm. (I, K) Dorsal, up. **(M-N)** Percent change in apical cell area (M) and cell height (N) 40 minutes after wounding with respect to pre-wound values in embryos injected with DMSO (blue, *n* = 27 cells in 8 embryos in M, 36 cells in 6 embryos in N), or rapamycin (red, *n* = 30 cells in 7 embryos in M, 36 cells in 6 embryos in N). (C, E) Error bars, s.e.m.. (D, F-H, M-N) Error bars, standard deviation, s.d.; boxes, s.e.m.; gray lines, mean. n.s., not significant; * *P* < 0.05.

### mTORC1 is dispensable for the molecular rearrangements that drive rapid wound healing

To investigate the mechanisms by which mTORC1 promotes rapid wound repair, we examined whether mTOR inhibition disrupted the cytoskeletal rearrangements associated with the wound healing response. Rapamycin treatment did not affect myosin polarization or dynamics around the wound (Figure 1A-B). Myosin levels at the wound margin increased by 2.5 ± 0.5-fold in controls (mean ± standard error of the mean, s.e.m.) and by 2.3 ± 0.3-fold in rapamycin-injected embryos (Figure 1E). The rate of myosin accumulation around the wound, measured as the half-time of myosin fluorescence, was not affected by mTOR inhibition (15 ± 6 min in controls *vs.* 15 ± 4 min in rapamycin-treated embryos, Figure 1F). We obtained similar results when we measured myosin dynamics at the wound edge in *raptor* RNAi embryos (Figure S1E); or when we quantified F-actin dynamics around the wound in embryos expressing UtrophinABD:GFP as a marker of filamentous actin (Rauzi et al., 2010) and treated with rapamycin or with DMSO as a control (Figure S2A-D). Thus, mTORC1 does not affect cytoskeletal rearrangements during embryonic wound closure.

Adherens junction remodelling is a pre-requisite for the cytoskeletal polarization that drives rapid wound healing (Hunter et al., 2015; Matsubayashi et al., 2015). Specifically, adherens junctions are depleted via endocytosis from bicellular contacts between wounded cells and cells adjacent to the wound, and accumulate at former tricellular contacts at the wound edge (Brock et al., 1996; Wood et al., 2002; Abreu-Blanco et al., 2012; Zulueta-Coarasa et al., 2014; Hunter et al., 2015; Matsubayashi et al., 2015; Rothenberg et al., 2023). We thus quantified E-cadherin levels at the wound edge in DMSO-injected controls and in rapamycin-treated embryos. Adherens junction remodelling was not affected by mTOR inhibition: E-cadherin levels at bicellular contacts at the wound edge decreased by 38 ± 9% in DMSO-injected embryos 15 min after wounding, and by 38 ± 8% in embryos treated with rapamycin (Figure S2E-I). At former tricellular junctions, E-cadherin levels increased by 14± 8% in controls 15 min after wounding, and by 16 ± 7% in rapamycin-treated embryos (Figure S2E-H, J). Together, our data show that mTOR signalling is not necessary for the molecular rearrangements that drive rapid wound repair.

### mTORC1 restricts wound size by limiting cell damage

To understand how mTORC1 could be facilitating rapid wound healing, we examined the size of the wounds. We found that rapamycin treatment resulted in wounds that were 44% larger than in controls (*P* < 0.05, Figure 1C, G). Similarly, wounds in *raptor* RNAi embryos were 25% larger than in controls (*P* < 0.05, Figure S1C, F). As indicated above, the increased wound area when we disrupted mTORC1 signalling was not associated with defects in myosin polarization to the wound edge (Figures 1E and S1E), which can cause large wounds (Kobb et al., 2017). Furthermore, laser ablation experiments and mechanical modelling of the response to ablation revealed that mTOR inhibition with rapamycin did not affect epidermal tension (Figure S3A-C) or viscoelasticity (Figure S3D). Together, our results suggest that changes to cell mechanics are not responsible for the increased wound area when mTORC1 signalling is disrupted.

We previously showed that p38 signalling limits wound size by protecting the cells around the wound from wound-induced damage (Scepanovic et al., 2021b). Notably, rapamycin treatment was associated with a 3-fold increase in the number of cells that had not been directly targeted by the laser but were enclosed by the myosin cable (0.9 ± 0.3 cells in controls *vs.* 2.5 ± 0.7 cells in rapamycin-treated embryos, *P* < 0.05, Figure 1A-B, H). Embryos expressing *raptor* RNAi displayed a 5-fold increase in the number of cells not targeted by the laser within the wound with respect to *mCherry* RNAi controls (0.5 ± 0.3 cells in *mCherry* RNAi *vs.* 2.4 ± 0.4 cells in *raptor* RNAi embryos, *P* < 0.05, Figure S1A-B, G). Together, our results suggest that mTORC1 limits cell damage upon wounding, thus facilitating rapid tissue repair.

### mTORC1 promotes cell swelling to drive rapid wound closure

Rapid wound closure requires an increase in the volume of the cells adjacent to the wound (Tanner et al., 2009; Scepanovic et al., 2021b). To determine whether mTOR contributes to rapid wound healing by promoting an increase in the volume of the cells around the wound, we quantified changes in cross-sectional area and height for cells adjacent to wounds in control and rapamycin-treated embryos expressing the membrane marker Resille:GFP (Buszczak et al., 2007). mTOR inhibition resulted in apical area changes that were 65% smaller than in controls (*P* < 0.05, Figure 1I, K, M), with no significant effects on cell heights (*P* < 0.05, Figure 1J, L, N). These results suggest that mTOR signalling is required for the increase in cell volumes associated with rapid embryonic wound repair.

### mTORC1 inhibits autophagy during embryonic wound closure

How does mTORC1 promote cell swelling? mTORC1 can increase cell size through protein synthesis (Brunn et al., 1997; Hara et al., 1997; Gingras et al., 1999; Holz et al., 2005), but protein translation is not required for rapid multicellular wound repair (Scepanovic et al., 2021b). mTOR can also promote cell swelling in epithelial cells by inhibiting autophagy (Orhon et al., 2016). To investigate whether mTOR regulates autophagy in the embryonic epidermis, we measured the fluorescence of mCherry:Atg8a, an autophagosome marker (Nezis et al., 2010), in the presence and absence of mTOR signalling. mCherry:Atg8a formed punctate structures resembling autophagosomes (smaller) and autolysosomes (larger) in the embryonic epidermis (Figure 2A) (Mauvezin et al., 2014). We found that rapamycin treatment increased mCherry:Atg8a levels in the intact epidermis by 61 ± 49% 15 minutes after injection (*P* < 0.05, Figure 2A-D, K), suggesting that mTOR restricts autophagy in the *Drosophila* embryonic epidermis.

**Figure 2.**
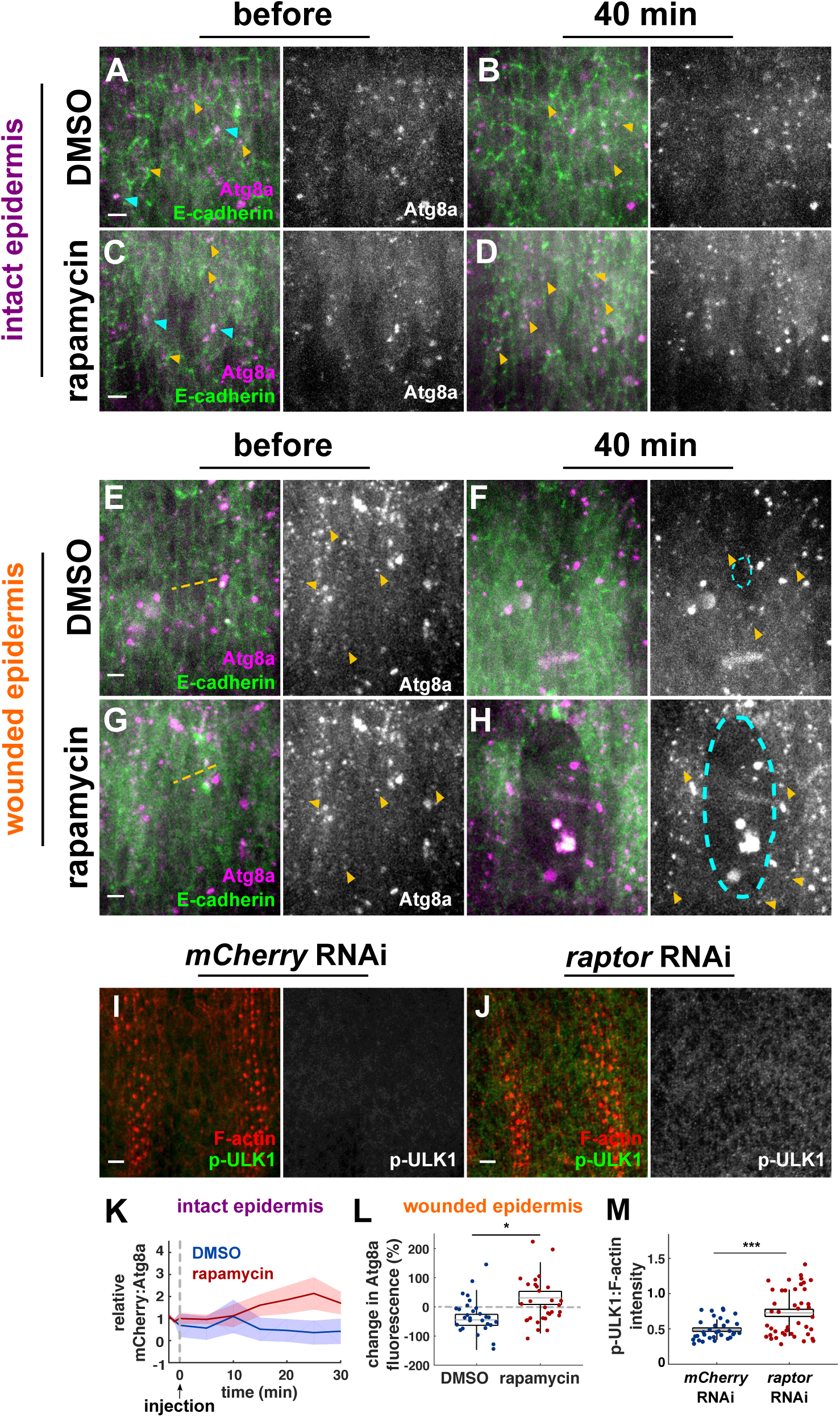
mTORC1 restricts autophagy during wound repair. **(A-H)** Epidermal cells in embryos expressing E-cadherin:GFP (green) and mCherry:Atg8a (magenta, grayscale), and injected with DMSO (A-B, E-F) or rapamycin (C-D, G-H). Yellow arrowheads indicate autophagosomes, blue arrowheads indicate autolysosomes, yellow dashed lines indicate wound sites, cyan dashed lines outline wounds. Time after injections (A-D) or wounding (E-H) is shown. **(I-J)** Epidermal cells in embryos expressing *mCherry* RNAi (I) or *raptor* RNAi (J) and immunostained for p-ULK1 (green, grayscale) and phalloidin to label F-actin (red). **(K-L)** Relative mCherry:Atg8a fluorescence in embryos injected with DMSO (blue, *n* = 39 cells in 5 embryos), or rapamycin (red, *n* = 38 cells in 4 embryos) (K), percent change in mCherry:Atg8a levels 40 min after wounding in cells adjacent to the wound in embryos injected with DMSO (blue, *n* = 30 cells in 5 embryos), or rapamycin (red, *n* = 30 cells in 6 embryos) (L), and ratio of p-ULK1 intensity inside epidermal cells to cortical F-actin levels in embryos expressing *mCherry* RNAi (blue, *n* = 36 cells in 5 embryos) and *raptor* RNAi (red, *n* = 43 cells in 6 embryos) (M). (A-J) Anterior left, dorsal up. Bars, 5 μm. (K) Error bars, s.e.m.. (L-M) Error bars, s.d.; boxes, s.e.m.; gray lines, mean. * *P* < 0.05, *** *P* < 0.001.

We investigated whether autophagy levels must be limited for rapid wound closure. To quantify autophagy in wounded embryos, we measured mCherry:Atg8a fluorescence in cells surrounding epidermal wounds (Figure 2E-F, Video S2). We found that mCherry:Atg8a fluorescence decreased by 44 ± 20% in the cells surrounding the wound 40 minutes after wounding (*P* < 0.0001, Figure 2L), suggesting that downregulating autophagy may be an important step for rapid wound repair. To determine whether mTORC1 limits autophagy during embryonic wound closure, we measured mCherry:Atg8a fluorescence in the cells adjacent to embryonic wounds when mTOR was inhibited with rapamycin (Figure 2G-H). Rapamycin treatment prevented the loss of mCherry:Atg8a fluorescence, which in fact increased by 31 ± 22% during the course of wound repair (*P* < 0.0001, Figure 2L), demonstrating a significant effect of mTOR inhibition relative to controls (*P* < 0.05). We validated our results using a second marker for autophagosomes, GFP:Atg8a (Arsham and Neufeld, 2009), which only labels autophagosomes prior to lysosome fusion, as GFP fluorescence is quenched in the lysosome (Mauvezin et al., 2014) (Figure S4A-D, Video S3). Rapamycin treatment increased GFP:Atg8a fluorescence 40 minutes after wounding by 86% with respect to controls (*P* < 0.05, Figure S4E-F). Thus, mTOR signalling suppresses autophagy during embryonic wound repair.

mTORC1 inhibits autophagy *in vitro* by preventing activation of the autophagy initiation factor ULK1 (Kim et al., 2011). The serine/threonine kinase AMP-activated protein kinase (AMPK) phosphorylates and activates ULK1 at multiple sites, including the conserved Ser555, thus promoting autophagy (Shang et al., 2011; Yang et al., 2015). mTORC1 inhibits autophagy by phosphorylating ULK1 at Ser757 and disrupting the interaction between AMPK and ULKL1 (Kim et al., 2011). To determine whether mTORC1 inhibits ULK1 in *Drosophila* embryos, we fixed and immunostained embryos using an antibody against the AMPK-phosphorylated form of ULK1 (p-ULK1) (Venkatesh et al., 2016) (Figure 2I-J). We found that disrupting mTORC1 signalling in *raptor* RNAi embryos increased p-ULK1 levels by 47% with respect to *mCherry* RNAi controls (*P* < 0.001, Figure 2M), suggesting increased autophagosome formation. Together, our results suggest that mTORC1 prevents ULK1 activation in embryos to restrict autophagy.

### Autophagy is detrimental for rapid wound healing

To investigate the role of autophagy in embryonic wound repair, we acutely increased autophagy during wound closure independent of mTOR signalling. We stimulated autophagy by injecting embryos with 500 μM of the inositol monophosphatase inhibitor, L-690,330, which reduces the levels of inositol 1,4,5-trisphosphate, a negative regulator of autophagy (Atack et al., 1993; Sarkar et al., 2005). L-690,330 treatment increased GFP:Atg8a fluorescence before wounding with respect to controls by 28% (Figure S5A, C, E, Video S4). Notably, inducing autophagy with L-690,330 prevented the wound-associated reduction in autophagy, which remained significantly higher than in controls (*P* < 0.001, Figure S5A-D, F). Thus, L-690,330 treatment is an effective way to promote autophagy during *Drosophila* embryonic wound repair.

To determine if reducing autophagy is necessary for rapid wound healing, we stimulated autophagy using L-690,330 and quantified the dynamics of wound repair (Figure 3A-B, Video S5). We found that in the presence of excessive autophagy, wounds closed 46% slower than in controls (*P* < 0.05, Figure 3C-D), suggesting that autophagy is detrimental for rapid wound healing. Consistent with previous findings in post-embryonic tissues (Kakanj et al., 2022), inducing autophagy did not cause any effects on myosin polarization to the wound edge (Figure 3E). We obtained similar results when we induced autophagy by overexpressing the *Drosophila* ULK1 ortholog, Atg1 (Braden and Neufeld, 2016) (Figure S6A-B). We found that Atg1 overexpression resulted in wounds that closed 30% slower than in controls (*P* < 0.05, Figure S6C-D), with no effects on myosin dynamics (Figure S6E). Together, our results show that autophagy levels must be reduced for embryonic wounds to heal rapidly.

**Figure 3.**
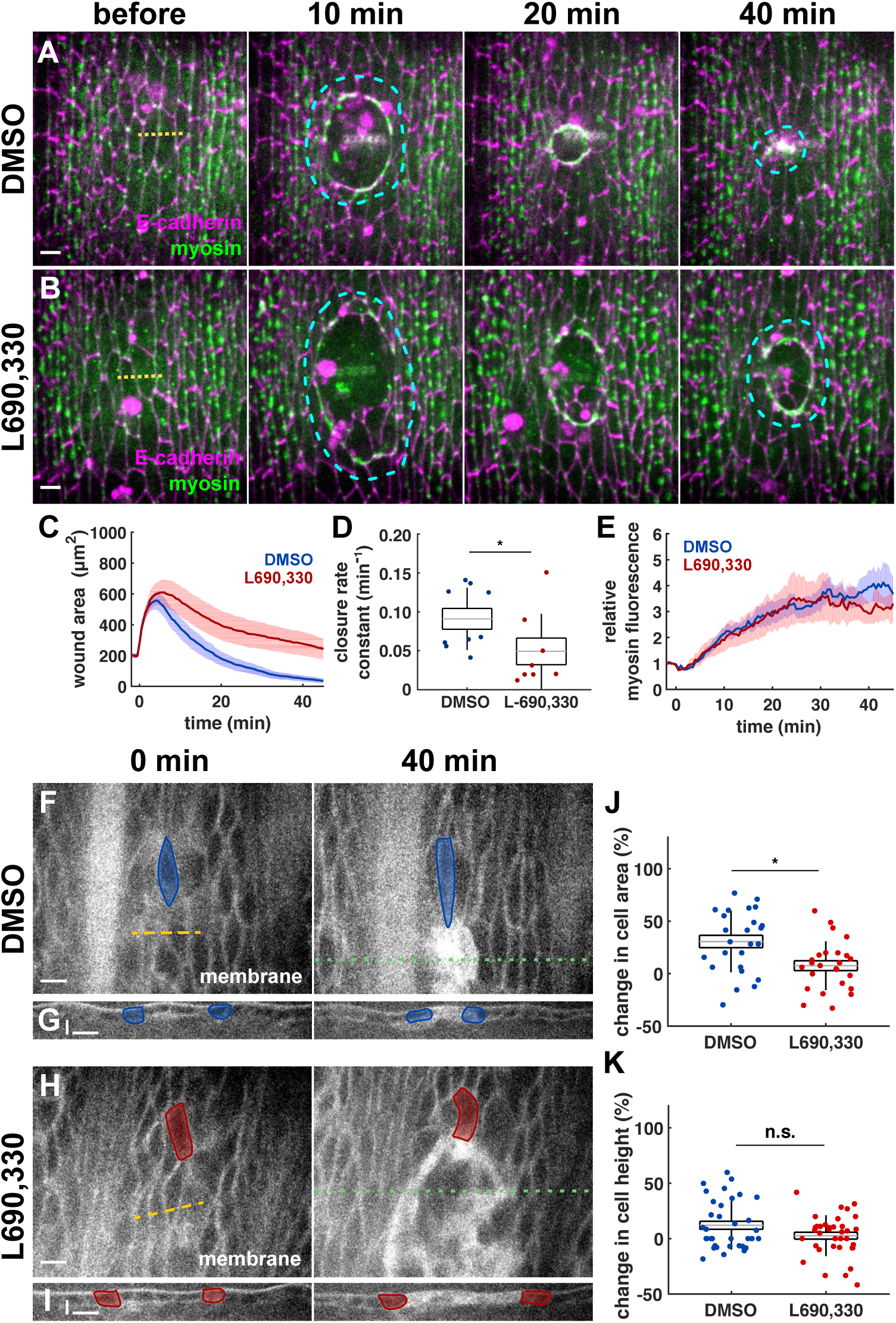
Autophagy slows down wound healing by inhibiting cell swelling. **(A-B)** Wound healing in embryos expressing myosin:GFP (Sqh, green) and E-cadherin:tdTomato (magenta), and injected with DMSO (A), or L690,330 (B). Yellow dashed lines indicate wound sites, cyan dashed lines outline wounds. Time after wounding is shown. **(C-E)** Wound area over time (C), wound closure rate constant (D), and relative mean myosin fluorescence at the wound edge over time (E) for embryos injected with DMSO (blue, *n* = 9 wounds), or L690,330 (red, *n* = 8). **(F-I)** Apical, XY (F, H), and cross-sectional, XZ (G, I) views of epidermal cells expressing Resille:GFP in embryos injected with 50% DMSO (F-G) or L-690,330 (H-I). Sample cells are highlighted in blue (DMSO) and red (L-690,330). Green dotted lines indicate the positions where cross-sections were taken. Time after injection is shown. **(J-K)** Percent change in apical cell area (J) and cell height (K) 40 minutes after wounding with respect to pre-wound values in embryos injected with DMSO (blue, *n* = 27 cells in 8 embryos in J, 36 cells in 6 embryos in K), or L-690,330 (red, *n* = 30 cells in 7 embryos in J, 36 cells in 6 embryos in K). (A-B, F-I) Anterior left. Bars, 5 μm. (A-B, F, H) Dorsal up. (C, E) Error bars, s.e.m.. (D, J-K) Error bars, s.d.; boxes, s.e.m.; gray lines, mean. n.s., not significant; * *P* < 0.05.

### Reduced autophagy facilitates cell swelling during wound repair

How does autophagy disrupt wound repair? Excessive autophagy can disrupt adherens junctions in the *Drosophila* larval epidermis (Kakanj et al., 2022). We thus quantified E-cadherin levels at the wound edge in DMSO-injected controls and in embryos in which we induced autophagy with L-690,330 (Figure S7A-B). We found that E-cadherin levels at bicellular contacts between wounded and adjacent cells decreased by 33 ± 6% in controls 15 min after wounding, and by 32 ± 6% in embryos injected with L-690,330 (Figure S7C-E). In the same period of time, E-cadherin accumulated by 20 ± 9% at tricellular junctions around the wound in controls, and by 21 ± 12% in embryos with excessive autophagy (Figure S7A-F). These data suggest that excessive autophagy does not disrupt adherens junction remodelling during embryonic wound closure.

Autophagy can reduce epithelial cell volumes (Orhon et al., 2016), and cell swelling is required for rapid wound closure (Tanner et al., 2009; Kennard and Theriot, 2020; Scepanovic et al., 2021b). Thus, we investigated whether autophagy disrupted cell volume increases during embryonic wound repair. We quantified changes in the cross-sectional area and the height of the cells adjacent to the wound in embryos expressing Resille:GFP, and injected with DMSO or with L-690,330 (Figure 3F-I). In control embryos, the cross-sectional area and the height of the cells adjacent to the wound increased significantly 40 minutes after wounding (31 ± 6% and 12 ± 4 %, respectively, Figure 3F-G, J-K). In contrast, when we treated embryos with L-690,330, the change in cross-sectional area associated with wound closure was 74% lower than in controls (8 ± 5%, *P* < 0.05, Figure 3H-I, J-K), with no significant change in cell heights. Together, our results suggest that autophagy disrupts rapid wound healing by preventing cell swelling.

The mechanisms by which autophagy regulates cell volume are not understood. Reduced autophagy is associated with cell swelling in various cells and tissues, including embryonic fibroblasts (Jin et al., 2015), kidney (Kimura et al., 2011; Orhon et al., 2016), liver (Komatsu et al., 2005; Jin et al., 2015) and intestinal cells (Chang et al., 2013). Autophagy diverts energy away from mass accumulation and can degrade mitochondria (Neufeld, 2012; Onishi et al., 2021). Given that cell mass and mitochondrial content must scale with cell volume (Kitami et al., 2012; Miettinen and Bjorklund, 2016; Demian et al., 2019), autophagy may counter cell swelling. Autophagy may also reduce the pool of plasma membrane available to accommodate an increase in cell volume, either by using up membrane to form autophagosomes (Ravikumar et al., 2010) or by degrading the trafficking machinery necessary to remodel the membrane, thus resulting in slower wound repair.

### mTORC1 inhibits autophagy to promote rapid wound healing

We reasoned that if mTORC1 inhibits autophagy to drive rapid wound repair, then reducing autophagy should rescue the wound closure defects associated with mTOR inhibition. To address that possibility, we reduced autophagy by expressing *atg8a* RNAi. *atg8a* RNAi did not significantly affect wound size, the rate of wound closure, or myosin polarization to the wound edge (Figure 4A, C, E-H, Video S6), consistent with previous data that autophagy is dispensable for post-embryonic wound healing (Kakanj et al., 2022). Strikingly, knocking down Atg8a in rapamycin-treated embryos increased the rate of wound closure by 60% with respect to rapamycin-treated embryos expressing *mCherry* RNAi (Figure 4A-G), with no significant effects on myosin dynamics (Figure 4H). We obtained similar results by acutely reducing autophagy using bafilomycin A1 (bafilomycin), which blocks autophagosome-lysosome fusion (Mauvezin and Neufeld, 2015). Treatment with 250 μM of bafilomycin with rapamycin increased the rate of wound closure by 2-fold with respect to rapamycin treatment (*P* < 0.05), to a level comparable to DMSO-injected controls (Figure S8A-B, D-G). Together, our results indicate that mTORC1 limits autophagy to drive rapid embryonic wound closure.

**Figure 4.**
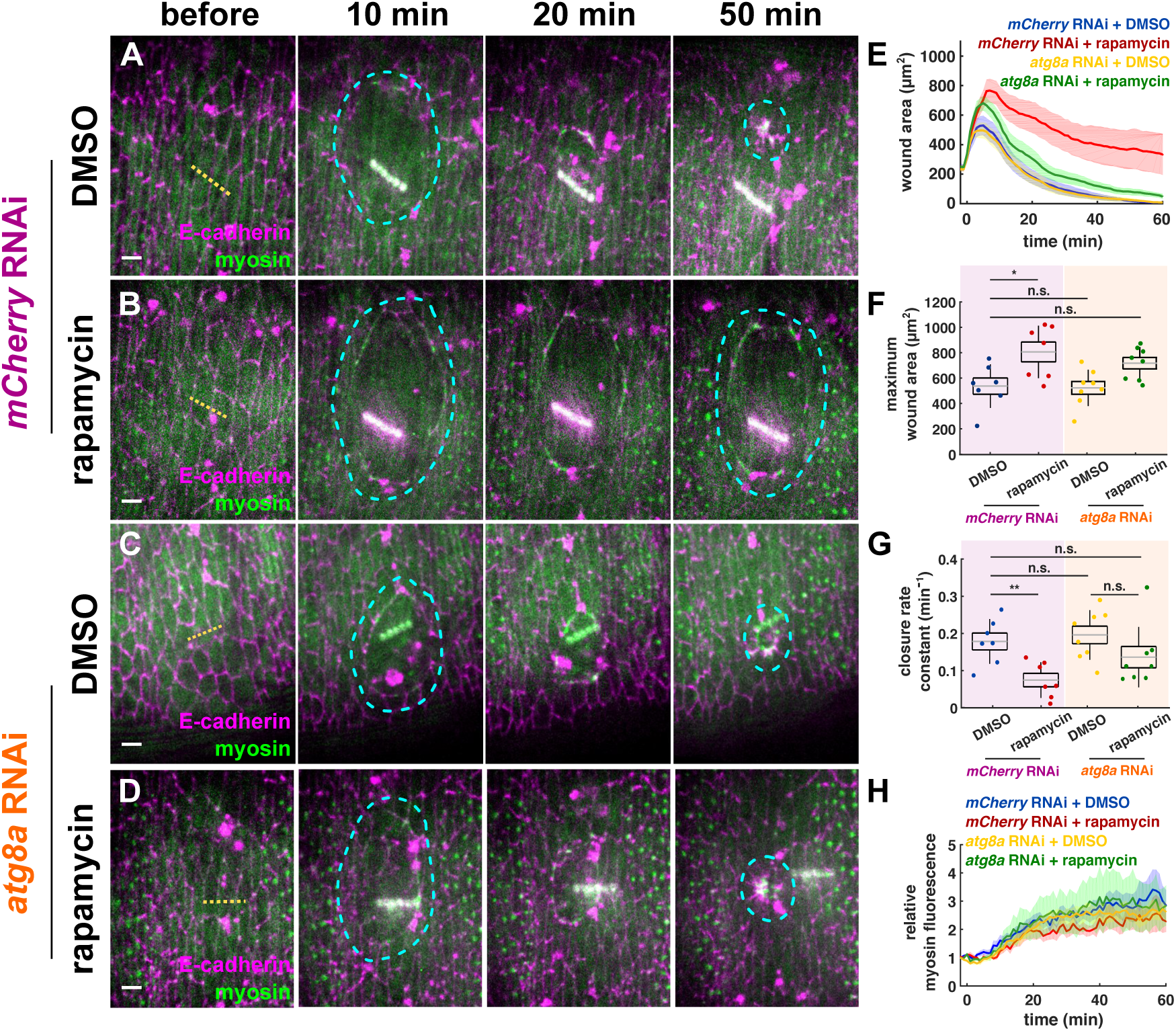
Inhibiting autophagy when mTOR signalling is disrupted rescues rapid wound healing. **(A-D)** Wound closure in embryos expressing myosin:GFP (Sqh, green) and E-cadherin:tdTomato (magenta), and *mCherry* RNAi (A-B) or *atg8a* RNAi (C-D). Embryos were injected with DMSO (A, C) or rapamycin (B, D). Yellow dashed lines indicate wound sites, cyan dashed lines outline wounds. Time after wounding is shown. Anterior left, dorsal up. Bars, 5 μm. **(E-H)** Wound area over time (E), maximum wound area (F), wound closure rate constant (G), and relative mean myosin fluorescence at the wound edge (H) for embryos expressing *mCherry* RNAi and injected with DMSO (blue, *n* = 7 wounds) or rapamycin (red, *n* = 7), or expressing *atg8a* RNAi and injected with DMSO (yellow, *n* = 8), or rapamycin (green, *n* = 8). (E, H) Error bars, s.e.m.. (F-G) Error bars, s.d.; boxes, s.e.m.; gray lines, mean. n.s., not significant; * *P* < 0.05, ** *P* < 0.01.

How is mTOR activated upon wounding? In mammalian epithelial cells and cardiomyocytes, phosphorylation of the mTORC1 downstream effector, S6K1, is dependent on oxidative stress (Sarbassov and Sabatini, 2005; Oka et al., 2017). Two conserved components of the mTORC1 complex, mTOR and MLST-8, harbour oxidant-modifiable cysteines (Meng et al., 2021), which could mediate mTORC1 activation in response to wounding. The PTEN phosphatase can also regulate mTORC1 by inhibiting the PI3K/AKT signalling pathway (Ghosh et al., 2013; Porta et al., 2014). Oxidative stress can inhibit PTEN, resulting in the enhanced activation of the serine/threonine kinase Akt1 (Leslie et al., 2003) an upstream activator of mTORC1 (Gingras et al., 1998; Miron et al., 2001; Peng et al., 2003). Consistent with this model, PTEN overexpression disrupts embryonic wound healing (Pickering et al., 2013).

Overall, our data show that autophagy is strictly regulated during embryonic wound healing. We propose that mTORC1 inhibits autophagy to facilitate cell swelling and drive rapid embryonic tissue repair. Our data highlight the importance of controlling autophagy for wound closure to proceed rapidly, and reveal new molecular targets that may enable pharmacological interventions to promote tissue repair when it is impaired, for instance, in diabetic patients with chronic skin wounds, in which autophagy is excessive (Guo et al., 2016).

## ACKNOWLEDGEMENTS

We are grateful to Tony Harris, Ran Kafri, Ashley Bruce, Negar Balaghi, Katy Rothenberg, Ana Maria do Carmo and Veronica Castle for comments on the manuscript, Michael Galko and Jonathan Palozzi for useful discussions, and Thomas Hurd and Gábor Juhász for reagents and flies. Flybase provided important information for this study. GS was supported by an Ontario Graduate Scholarship and an Alexander Graham Bell Canada Graduate Scholarship from the Natural Sciences and Engineering Research Council of Canada (NSERC). Our research is funded by grants to RFG from the Canadian Institutes of Health Research (156279 and 186188), NSERC (418-438-13), the Canada Foundation for Innovation (30279), and the Translational Biology and Engineering Program from the Ted Rogers Centre for Heart Research. RFG is the Canada Research Chair in Quantitative Cell Biology and Morphogenesis.

## AUTHOR CONTRIBUTIONS

GS – conceptualization, methodology, validation, formal analysis, investigation, data curation, writing (original draft, review and editing), visualization, funding acquisition; RFG – conceptualization, software, resources, writing (original draft, review and editing), supervision, project administration, funding acquisition.

## SUPPLEMENTAL VIDEO LEGENDS

**Video S1. mTOR signalling is necessary for rapid embryonic wound closure.**

Wound closure in embryos expressing myosin:GFP (Sqh, green) and E-cadherin:tdTomato (magenta) and injected with DMSO (left) or rapamycin (right). Images were acquired every 30 s. Time after wounding is shown. Anterior left, dorsal up. Bar, 5 μm.

**Video S2. mCherry:Atg8a levels are regulated in wounded embryos.**

Wound repair in embryos expressing mCherry:Atg8a and injected with DMSO (left) or rapamycin (right). Images were acquired every 60 s. Time after wounding is shown. Anterior left, dorsal up. Bar, 5 μm.

**Video S3. mTORC1 restricts autophagy during embryonic wound repair.**

Wound healing in embryos expressing GFP:Atg8a and injected with DMSO (left) or rapamycin (right). Images were acquired every 30 s. Time after wounding is shown. Anterior left, dorsal up. Bar, 5 μm.

**Video S4. L-690,330 induces autophagy during *Drosophila* embryonic wound healing.**

Wounded embryos expressing GFP:Atg8a and injected with DMSO (left) or L-690,330 (right). Images were acquired every 30 s. Time after wounding is shown. Anterior left, dorsal up. Bar, 5 μm.

**Video S5. Increased autophagy slows down wound closure.**

Wound healing in embryos expressing myosin:GFP (Sqh, green) and E-cadherin:tdTomato (magenta) and injected with DMSO (left) or L-690,330 (right). Images were acquired every 30 s. Time after wounding is shown. Anterior left, dorsal up. Bars, 5 μm.

**Video S6. Inhibiting autophagy rescues rapid wound repair in embryos with disrupted mTOR signalling.**

Wound healing in *mCherry* RNAi embryos injected with DMSO (left) or rapamycin (center-left), and in embryos expressing *atg8a* RNAi and injected with DMSO (center-right) or rapamycin (right). Embryos expressed myosin:GFP (Sqh, green) and E-cadherin:tdTomato (magenta).

Images were acquired every 60 s. Time after wounding is shown. Anterior left, dorsal up. Bar, 5 μm.

## MATERIAL AND METHODS

### Fly husbandry

All fly stocks were kept at 18°C or 25°C, on standard medium (cornmeal, yeast, agar, and molasses) provided by a central kitchen operated by H. Lipshitz. Stage 14-15 embryos (12-14 hours after egg laying) were collected overnight from apple juice-agar plates in mesh cages at 25°C. All experiments were completed within two weeks to minimize environmental changes (temperature, humidity, etc.) that could affect the results (Kuntz and Eisen, 2014).

We used the following transgenic lines for live imaging: *endo-DE-cadherin:tdTomato* (BDSC #58789) (Huang et al., 2009a), *sqh–sqh:GFP* (BDSC #57145) (Royou et al., 2003), *sqh-utrophinABD:GFP* (Rauzi et al., 2010), and *resille:GFP* (Buszczak et al., 2007). *daughterless*-*Gal4* (Perrin et al., 2003) was used to drive UAS transgenes. We used *UAS-raptor* RNAi (Perkins et al., 2015) to knockdown mTORC1 signalling, *UAS-atg8a* RNAi (BDSC# 34340) to reduce autophagy, and *UAS-mCherry* RNAi (BDSC# 35785) as controls. *UAS-atg1^6B^* flies (BDSC# 51655) (Mohseni et al., 2009) were used to induce autophagy, with *yw* flies crossed to the Gal4 driver as controls. To visualize autophagosomes we used *UAS-mCherry-atg8a* (Chang and Neufeld, 2009) and *UAS-GFP-atg8a* (BDSC# 51656). *endo-DE-cadherin:tdTomato* (Huang et al., 2009a) flies were used for fixation and staining.

### Embryo mounting and drug treatments

*Drosophila* embryos were prepared as described before (Scepanovic et al., 2021a). Briefly, embryos were dechorionated in 50% bleach, staged and transferred onto a coverslip. To prepare for injection, embryos were dehydrated for 7-11 minutes in a container full of silica beads (Drierite), then covered with a 1:1 mix of halocarbon oil 27 and 700 (Sigma-Aldrich).

Drug injections were done using a Transferman NK2 micromanipulator (Eppendorf) and a PV820 microinjector (World Precision Instruments) coupled to a spinning disk confocal microscope. Drugs were injected into the perivitelline space, where they are predicted to be diluted by 50-fold (Foe and Alberts, 1983). Drug injections were conducted immediately before wounding unless stated otherwise. All drugs were diluted in 50% DMSO. The following small molecule inhibitors were used: rapamycin (500 μM, BioShop), L-690,330 (500 μM, Sigma), and bafilomycin (250 μM, Cell Signaling Technology).

### Time-lapse imaging and laser ablation

All imaging was done at room temperature using a Revolution XD spinning disk confocal microscope (Andor Technology) with an iXon Ultra 897 camera (Andor Technology), 60X (NA 1.35) or 100X (NA 1.40) oil-immersion lenses (Olympus), and Metamorph software (Molecular Devices). Sixteen-bit Z-stacks were acquired every 30 seconds at 0.5 µm steps, or every 60 seconds at 0.3 µm steps, unless stated otherwise. For retraction velocity measurements, embryos were imaged every 4 seconds at 0.5 µm steps. Maximum intensity projections were used for all analyses.

A Micropoint nitrogen laser (Andor Technology) tuned to 365 nm was used to administer laser ablations. We delivered 10 pulses at discrete spots 2-μm apart along a 12-μm line on the epidermis to produce a multicellular wound. For spot laser ablations, 10 pulses were delivered at a single point over the course of 670 ms. Time 0 indicates the time of ablation for all experiments.

### Embryo fixation and staining

To prepare for immunostaining, embryos expressing *endo-DE-cadherin:tdTomato* were mounted as described above for injection. Embryos were washed off the coverslip using heptane, and fixed for 30 min in a 1:1 mix of heptane and 8% formaldehyde in phosphate buffer. Embryos were devitellinized by ethanol popping and stained with rabbit anti-phospho-ULK1 (Ser555) (1:50, Cell Signalling Technology) and goat anti-rabbit Alexa 488 (1:500, Invitrogen); and with phalloidin Alexa 647 (1:500, Invitrogen). Embryos were mounted in Prolong Gold (Molecular Probes) between two coverslips.

### Quantitative image analysis

Image analysis was performed using SIESTA (Fernandez-Gonzalez and Zallen, 2011; Leung and Fernandez-Gonzalez, 2015) and PyJAMAS (Fernandez-Gonzalez et al., 2022), and custom scripts written in MATLAB (Mathworks) using the DIPImage toolbox (TU Delft) and Python. To analyze wound closure dynamics, we segmented the wound using the semi-automated LiveWire algorithm in SIESTA (Fernandez-Gonzalez and Zallen, 2013), or MEDUSA, an automated tool to delineate the wound margin based on active contours (Zulueta-Coarasa et al., 2014).

To quantify the rate of wound healing, we measured the wound closure rate constant by fitting wound areas from the time of maximum wound expansion using the following exponential:

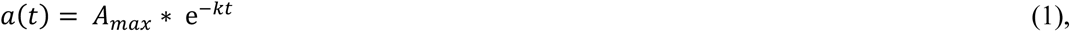

where *a*(*t*) represents the area of the wound at time *t* with respect to the time of maximum wound area, and *A_max_* and *k* are the two fitted parameters, with *A_max_* representing the maximum wound area, and *k* the wound closure rate constant.

To measure fluorescence at the wound margin, we quantified the mean pixel value under a 0.6-µm-wide mask over the wound edge. We normalized fluorescence intensities to their mean pre-wound values. Intensities were background subtracted using the image mode and corrected for photobleaching by dividing by the mean image intensity at each time point. The half-time of fluorescence accumulation was measured by fitting the following exponential to the fluorescence curves:

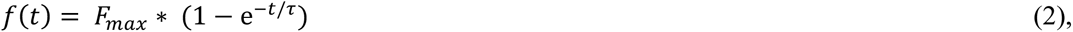

where *f*(*t*) is the mean fluorescence around the wound at time *t* with respect to the time of wounding, and *F_max_* and *τ* are two fitted parameters, with *F_max_* indicating the maximum fluorescence, and *τ* being a characteristic time scale related to the fluorescence half-time, *t*_1/2_, as:

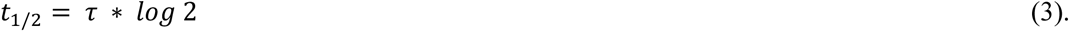

Adherens junction dynamics were quantified by selecting individual segments at the wound edge based on LiveWire annotations. Each segment was divided into 1000 evenly spaced points, and intensities were calculated using linear interpolation. Bicellular junction intensity was calculated using the mean of the central 300 points (Zulueta-Coarasa et al., 2014; Hunter et al., 2015). TCJ intensity was calculated by manually placing fiducials on the junctions and measuring the intensities from an extracted circular mask with a diameter of 0.5 μm (Rothenberg et al., 2023). Intensity values were normalized for photobleaching dividing by the mean image intensity.

To measure retraction velocity after laser ablation, the positions of the two tricellular vertices connected by the ablated junction were manually tracked. The instantaneous recoil velocity was quantified using the change in distance between the two vertices divided by the duration of the ablation plus the time necessary to acquire a stack. The mechanical properties of the severed junction were measured using a Kelvin-Voigt model consisting of a spring (elasticity) and dashpot (viscosity) configured in parallel (Kumar et al., 2006; Fernandez-Gonzalez et al., 2009). The model can be used to fit the distance retracted by the ends of the severed structure, *L*(*t*) at time *t* after ablation, using the following equation:

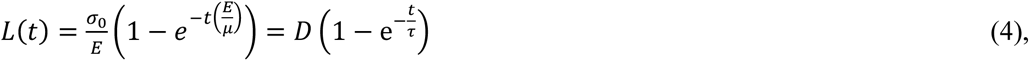

where σ_0_ is the tension sustained by the junction, *E* is the elastic modulus, and *µ* is the viscosity. *D* is the asymptotic distance retracted by the vertices, proportional to the force-to-elastic modulus ratio; and *τ* is a relaxation time proportional to the viscosity-to-elastic modulus ratio.

Cell volumes were estimated as described before (Scepanovic et al., 2021b). Briefly, the apical areas and heights of cells adjacent to the wound were measured in embryos expressing Resille:GFP. For area analysis, three apical slices were selected, projected and segmented using the LiveWire algorithm. To measure cell heights, we projected the stacks to obtain XZ views using ImageJ (Schneider et al., 2012) and manually selected 6 points around the wound edge for height measurement.

We estimated changes in autophagy in embryos expressing mCherry:Atg8a, by measuring the fluorescence in the pixels inside cell membrane annotations. To calculate the changes in autophagy in embryos expressing GFP:Atg8a, we used the Livewire wound edge segmentation. To restrict our measurements to cells adjacent to the wound, the wound edge segmentation was filled and subsequently dilated with a 7-μm-wide square structuring element; and we quantified the mean pixel values under a mask resulting from the difference between the dilated wound mask and the actual wound. For each wound, we quantified the change in fluorescence as the ratio between the average signal 40-41 minutes after wounding and the average pre-wound signal.

### Statistical analysis

Sample means were compared using a non-parametric Mann-Whitney test (Glantz, 2002). To compare more than two groups, we used a Kruskal–Wallis test to reject the null hypothesis, and Dunn’s test for pairwise comparisons. Paired samples were compared using a Wilcoxon Rank Sum Test.

---

**Figure S1.**
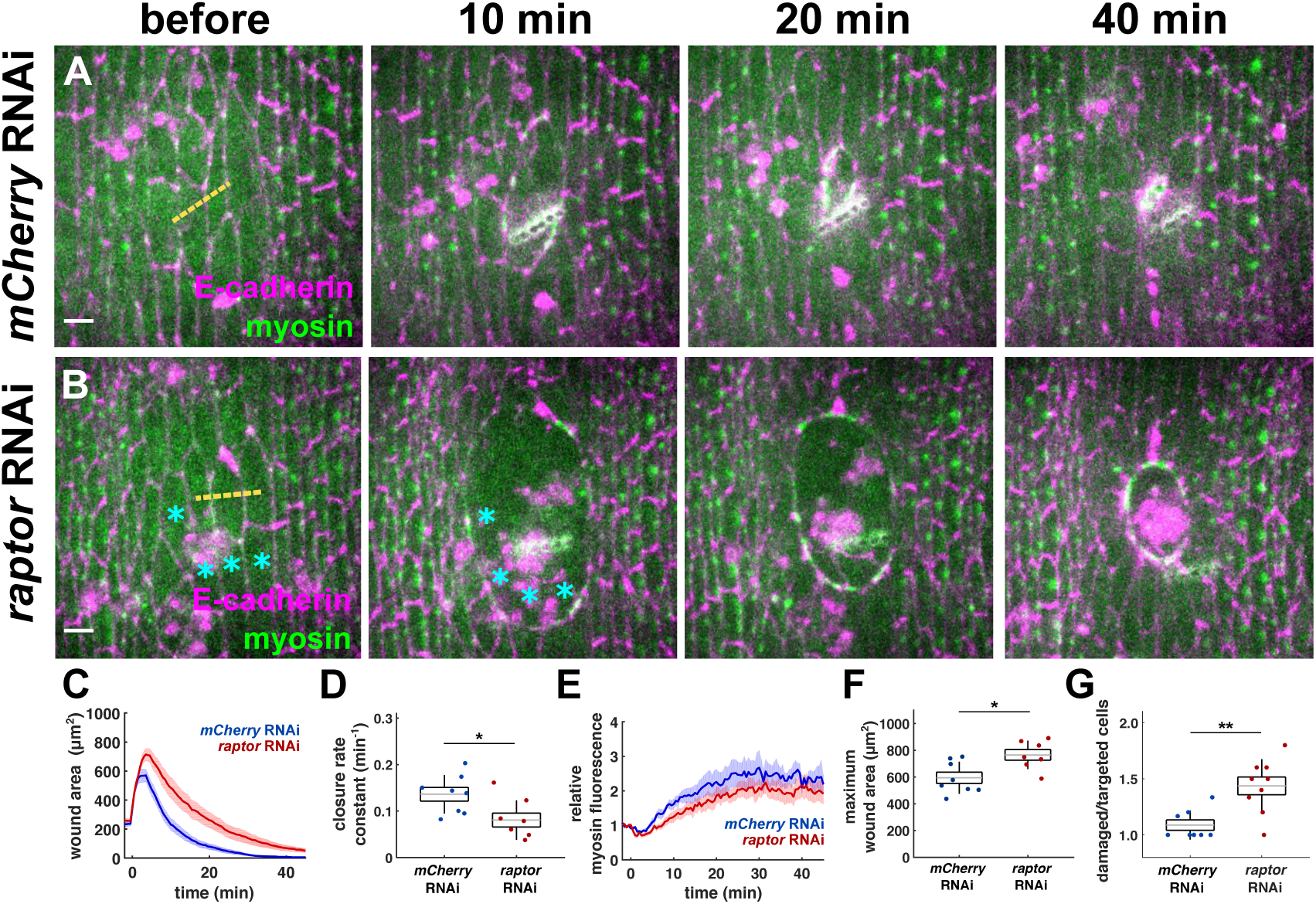
The mTORC1 component Raptor is required for rapid wound healing. **(A-B)** Wounded closure in embryos expressing myosin:GFP (Sqh, green) and E-cadherin:tdTomato (magenta), and *mCherry* RNAi (A) or *raptor* RNAi (B). Yellow dashed lines indicate wound sites. Cyan asterisks track cells not targeted by the laser but eventually included inside the wound. Time after wounding is shown. Anterior left, dorsal up. Bars, 5 μm. **(C-G)** Wound area over time (C), wound closure rate constant (D), relative mean myosin fluorescence at the wound edge over time (E), maximum wound area (F), and ratio of damaged cells (inside the myosin cable) to number of cells targeted by the laser (G) for embryos expressing *mCherry* RNAi (blue, *n* = 8 wounds), or *raptor* RNAi (red, *n* = 7, C-F, or *n* = 9, G). (C, E) Error bars, s.e.m.. (D, F-G) Error bars, s.d.; boxes, s.e.m.; gray lines, mean. * *P* < 0.05; ** *P* < 0.01.

**Figure S2.**
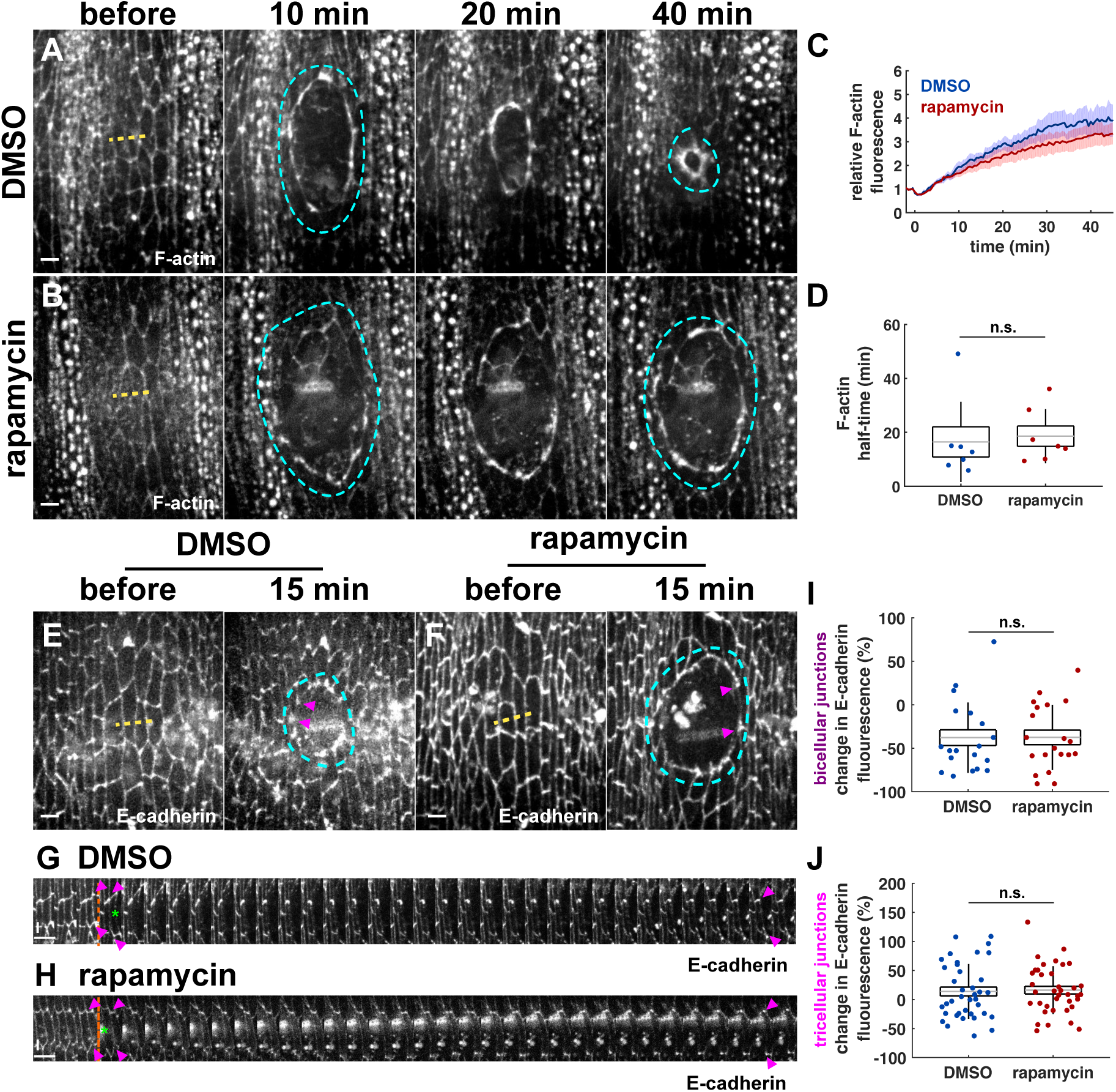
mTOR does not control F-actin or adherens junction dynamics during wound repair. **(A-B)** Wound repair in embryos expressing UtrophinABD:GFP and injected with DMSO (A), or rapamycin (B). Time after wounding is shown. **(C-D)** F-actin fluorescence at the wound edge over time (C) and half-time of F-actin fluorescence (D) for embryos injected with DMSO (blue, *n* = 7 wounds) or rapamycin (red, *n* = 7). **(E-F)** Epidermal cells in embryos injected with DMSO (E) or rapamycin (F) and expressing E-cadherin:tdTomato, before (left) and 15 minutes after wounding (right). **(G-H)** Kymographs showing the bicellular junctions at the wound edge indicated by magenta arrowheads in (E-F). Orange dashed lines show the time of ablation. Green asterisks indicate the position of the wound. Anterior left, dorsal up. Bars, 30 s - horizontal, 5 μm - vertical. **(I, J)** Percent change in E-cadherin:tdTomato levels 15 min after wounding in bicellular junctions (I) and in tricellular junctions (J) at the wound margin in embryos injected with DMSO (blue, *n* = 19 bicellular junctions in I and 38 tricellular junctions in J from 7 embryos) or rapamycin (red, *n* = 20 in I and 38 in J from 8 embryos). (A-B, E-F) Yellow dashed lines indicate wound sites, cyan dashed lines outline wounds. Anterior left, dorsal up. Bars, 5 μm. (C) Error bars, s.e.m.. (D, I-J) Error bars, s.d.; boxes, s.e.m.; gray lines, mean. n.s., not significant.

**Figure S3.**
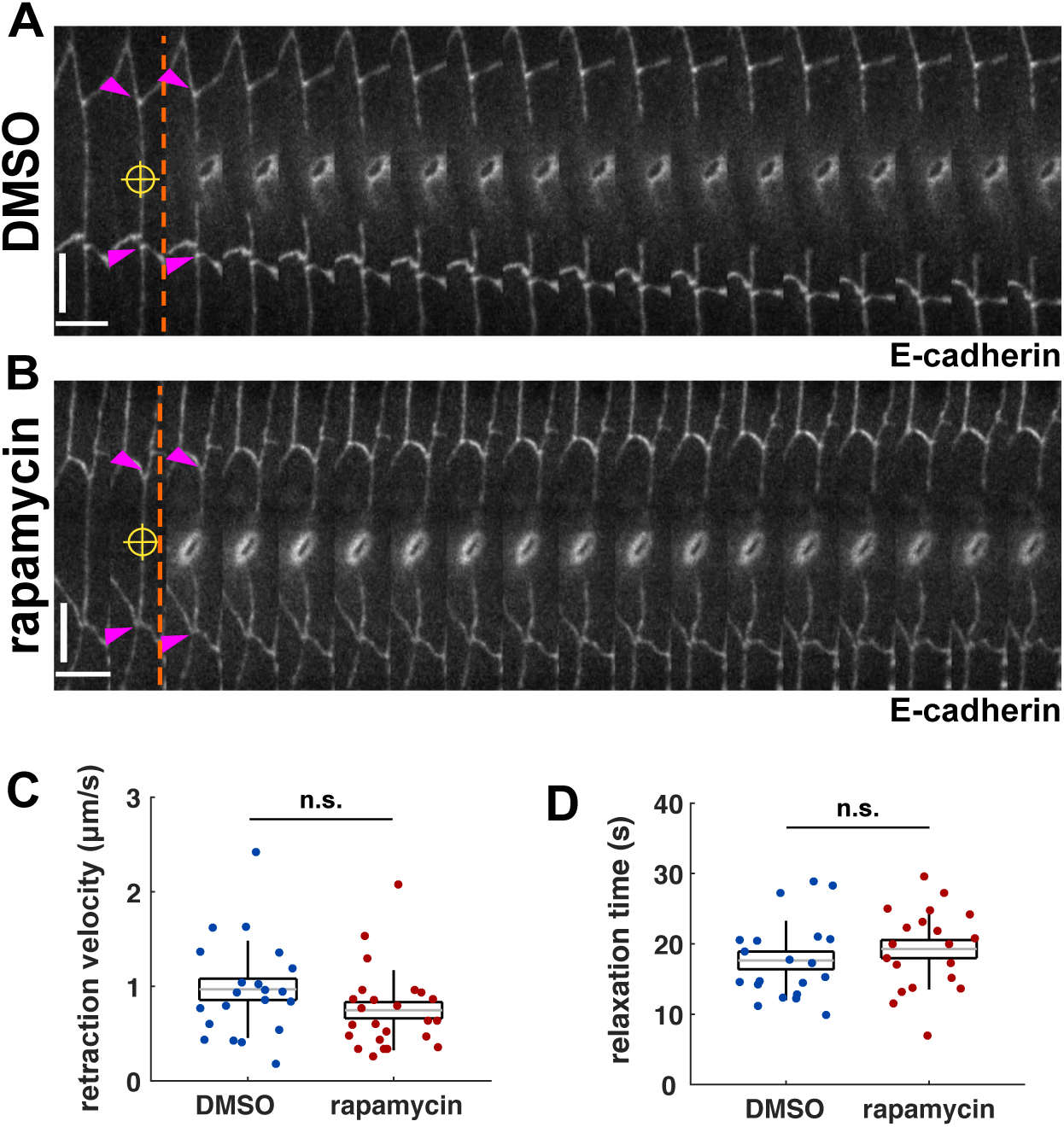
mTOR does not regulate cell mechanics. **(A-B)** Kymographs showing the recoil after laser ablation of a cell junction in embryos expressing Ecadherin:tdTomato and injected with DMSO (A) or rapamycin (B). Yellow targets denote the sites of ablation, orange dashed lines show the time of ablation. Magenta arrowheads indicate the vertices at the ends of the severed junctions, immediately before and after ablation. Anterior left, dorsal up. Bars, 4 s - horizontal, 5 μm - vertical. **(C-D)** Recoil velocity (C) and relaxation time (D) after laser ablation of cell junctions in embryos injected with DMSO (blue, *n* = 21 junctions in 21 embryos) or rapamycin (red, *n* = 24 junctions in 24 embryos). Error bars, s.d.; boxes, s.e.m.; gray lines, mean. n.s., not significant.

**Figure S4.**
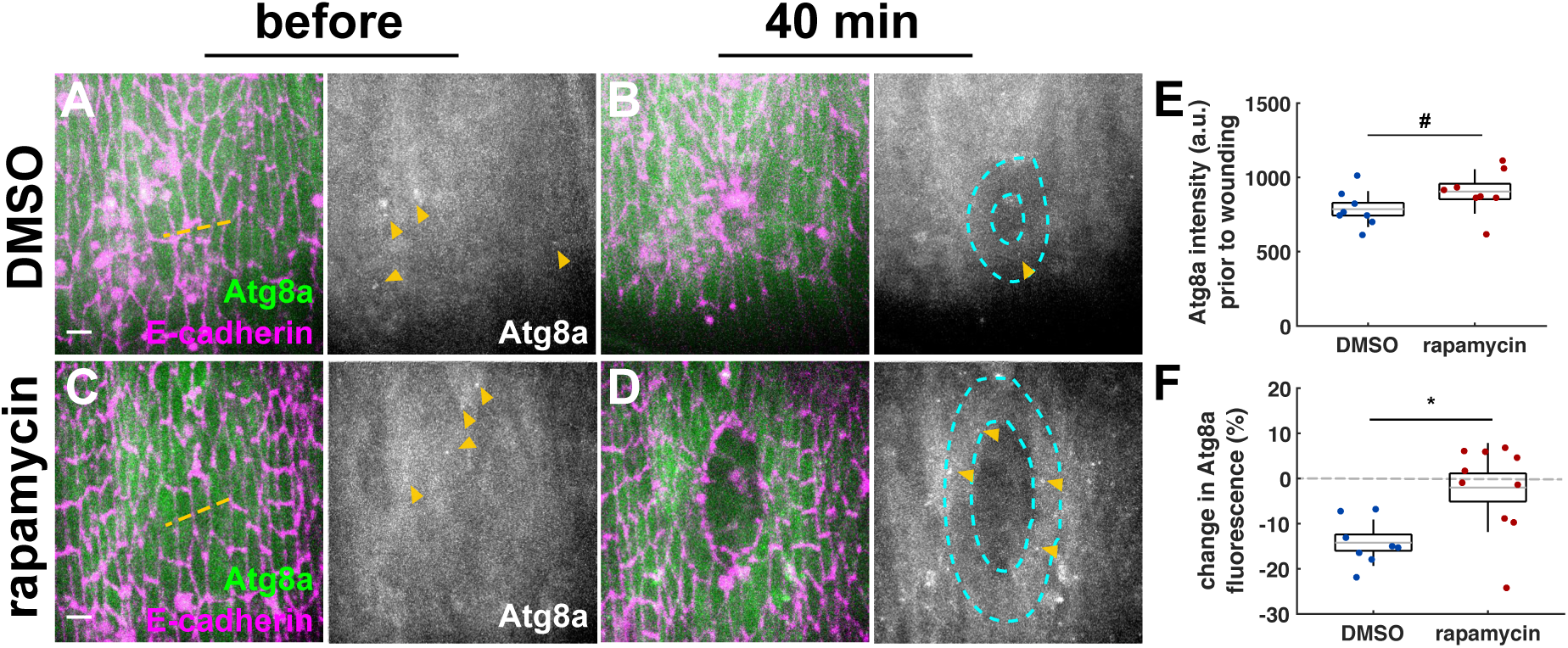
mTORC1 restricts autophagosome formation during embryonic wound repair. **(A-D)** Epidermal cells before (A, C) and 40 minutes after wounding (B, D) in embryos expressing E-cadherin:tdTomato (magenta) and GFP:Atg8a (green, grayscale). Embryos were injected with DMSO (A-B) or rapamycin (C-D). Yellow dashed lines indicate wound sites, cyan dashed lines outline wounds. Yellow arrowheads indicate autophagosomes. Time after wounding is shown. **(E-F)** GFP:Atg8a intensity prior to wounding measured 5 min after injection (E) and percent change in GFP:Atg8a levels 40 min after wounding (F) in cells adjacent to wounds in embryos treated with DMSO (blue, *n* = 8 wounds), or rapamycin (red, *n* = 10). Error bars, s.d.; boxes, s.e.m.; gray lines, mean. # *P* < 0.09, * *P* < 0.05.

**Figure S5.**
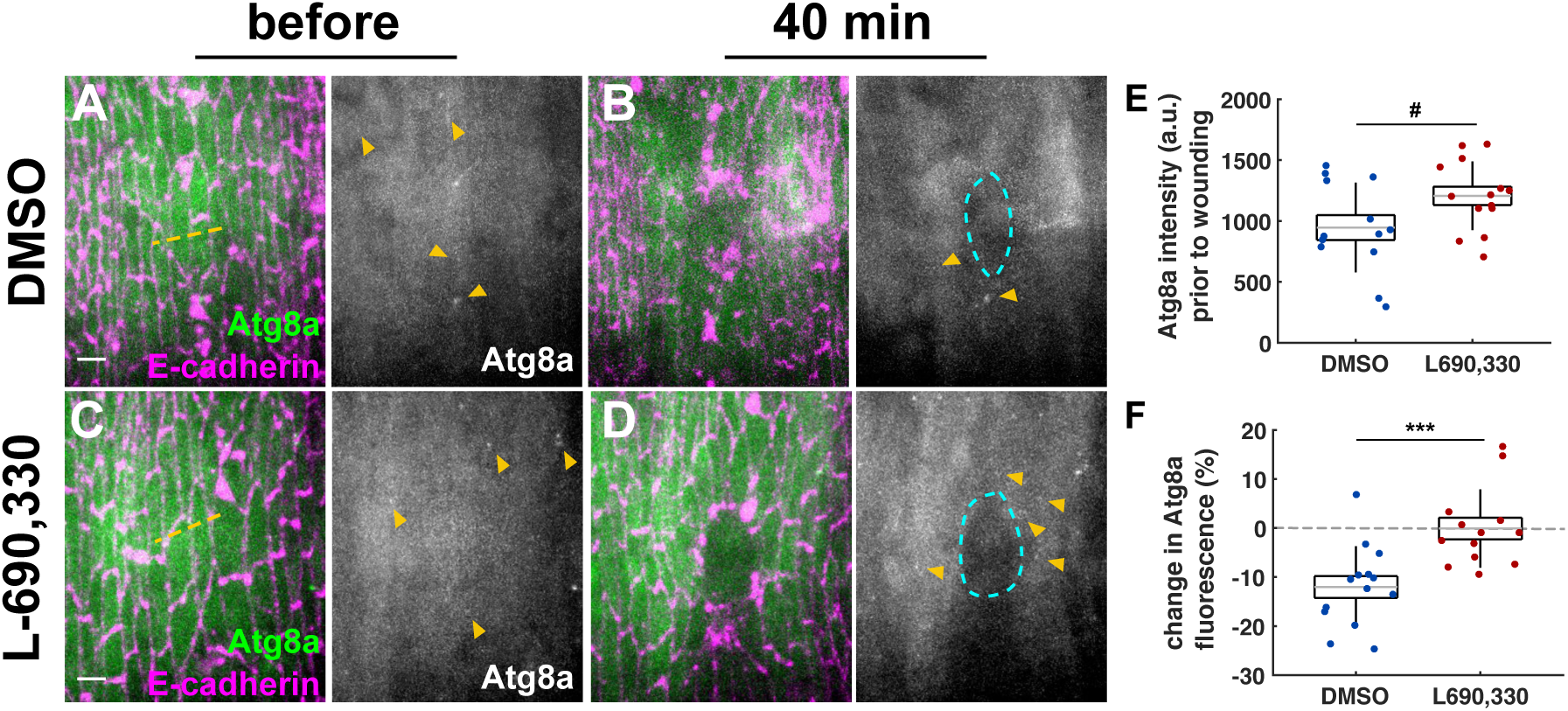
L-690,330 treatment prevents the reduction of autophagy associated with wound healing. **(A-D)** Epidermal cells before (A, C) and 40 minutes after wounding (B, D) in embryos expressing E-cadherin:tdTomato (magenta) and GFP:Atg8a (green, grayscale). Embryos were injected with DMSO (A-B) or the autophagy inducer L-690,330 (C-D). Yellow dashed lines indicate wound sites, cyan dashed lines outline wounds. Yellow arrowheads indicate autophagosomes. Time after wounding is shown. Anterior left, dorsal up. Bars, 5 μm. **(E-F)** GFP: Atg8a intensity prior to wounding measured 5 min after injection (E) and percent change in GFP:Atg8a levels 40 min after wounding (F) in cells adjacent to wounds in embryos treated with DMSO (blue, *n* = 14 wounds), or L-690,330 (red, *n* = 13). Error bars, s.d.; boxes, s.e.m.; gray lines, mean. # *P* < 0.1, *** *P* < 0.001.

**Figure S6.**
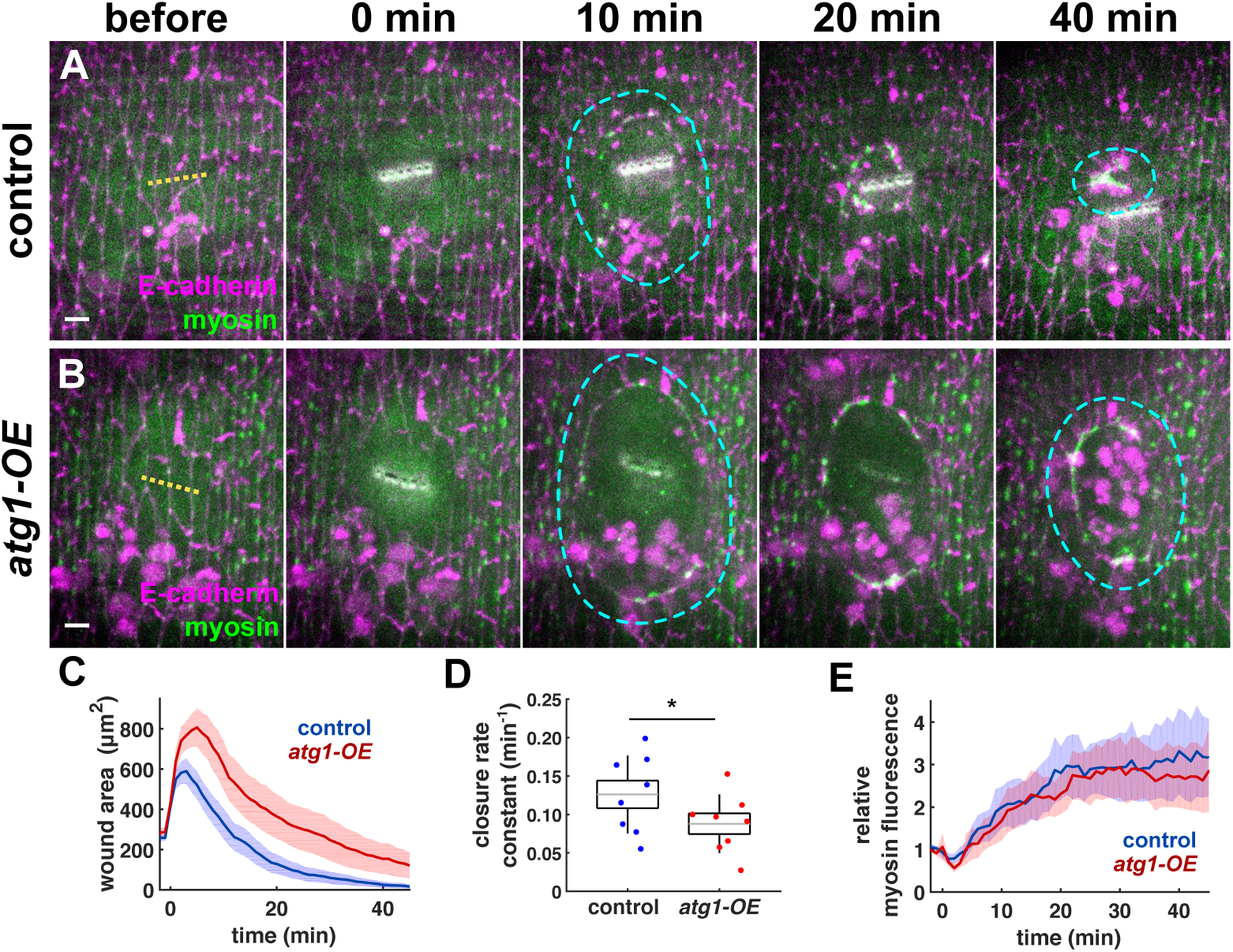
Atg1 overexpression disrupts rapid embryonic wound closure. **(A-B)** Wound closure in embryos expressing myosin:GFP (Sqh, green) and E-cadherin:tdTomato (magenta) in control embryos (A), or in embryos overexpressing Atg1 (*atg1-OE*) (B). Yellow dashed lines indicate wound sites, cyan dashed lines outline wounds. Time after wounding is shown. Anterior left, dorsal up. Bars, 5 μm. **(C-E)** Wound area over time (C), wound closure rate constant (D), and relative mean myosin fluorescence at the wound edge over time (E), for *yw* x *daughterless-Gal4* controls (blue, *n* = 8 wounds), or *UAS-atg1* x *daughterless-Gal4* Atg1- overexpressing embryos (red, *n* = 8). (C, E) Error bars, s.d.; boxes, s.e.m.; gray lines, mean. (D) Error bars, s.e.m.. * *P* < 0.05.

**Figure S7.**
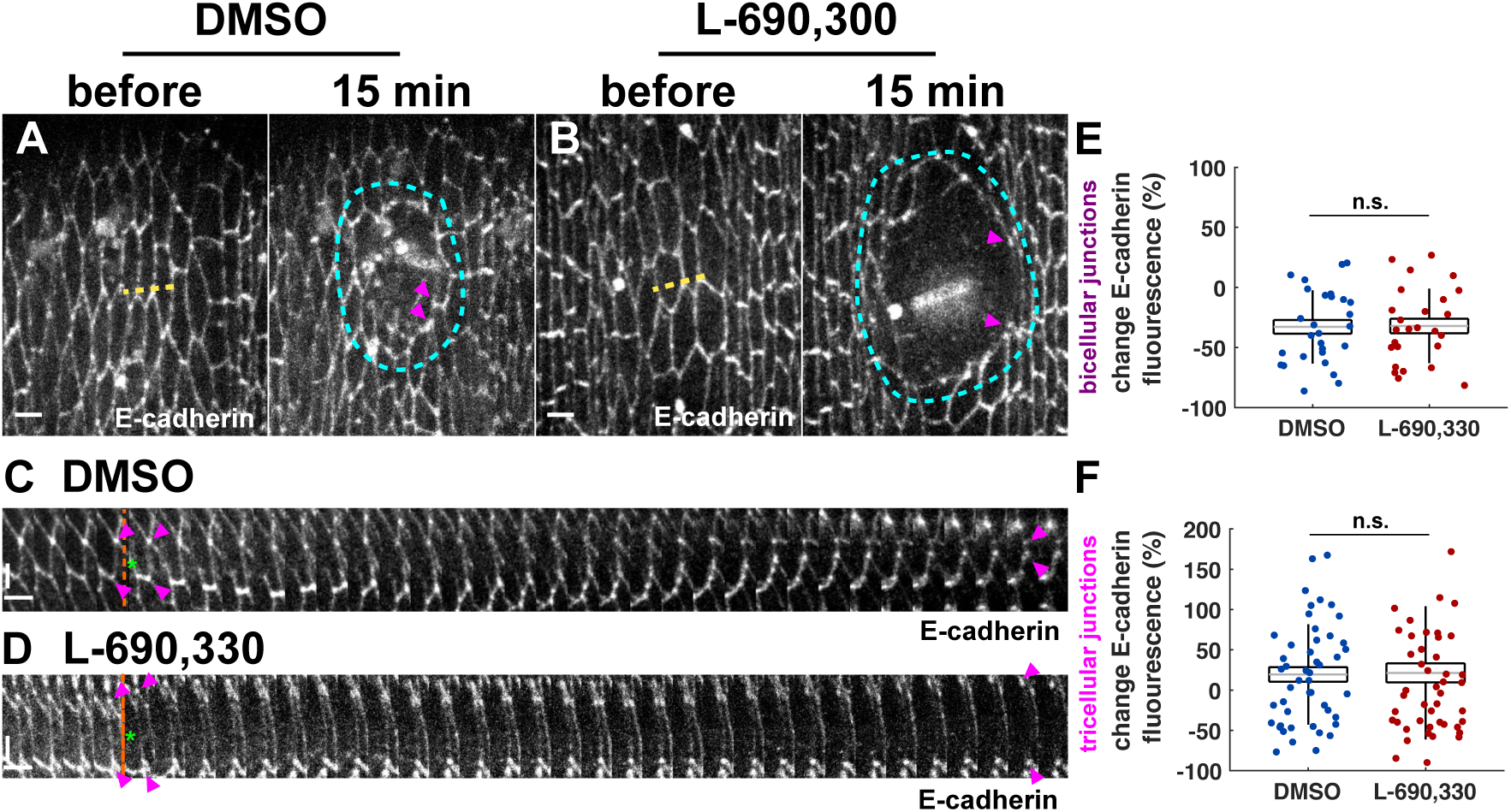
Autophagy is dispensable for adherens junction remodelling during wound repair. **(A-B)** Epidermal cells in embryos injected with DMSO (A) or L-690,330 (B) and expressing E-cadherin:tdTomato, before (left) and 15 minutes after wounding (right). Yellow dashed lines indicate wound sites, cyan dashed lines outline wounds. Bars, 5 μm. **(C-D)** Kymographs showing the bicellular junctions at the wound edge indicated by magenta arrowheads in (A-B). Orange dashed lines show the time of ablation. Green asterisks indicate the position of the wound. Bars, 30 s – horizontal, 5 μm - vertical. (A-D) Magenta arrowheads indicate the tricellular junctions flanking a bicellular contact at the wound margin. Anterior left, dorsal up. **(E, F)** Percent change in E-cadherin:tdTomato 15 min after wounding in bicellular junctions (E) and in tricellular junctions (F) at the wound margin in embryos injected with DMSO (blue, *n* = 28 bicellular junctions in E and 48 tricellular junctions in F from 9 embryos) or L-690,330 (red, *n* = 26 in E and 48 in F from 9 embryos). Error bars, s.d.; boxes, s.e.m.; gray lines, mean. n.s., not significant.

**Figure S8.**
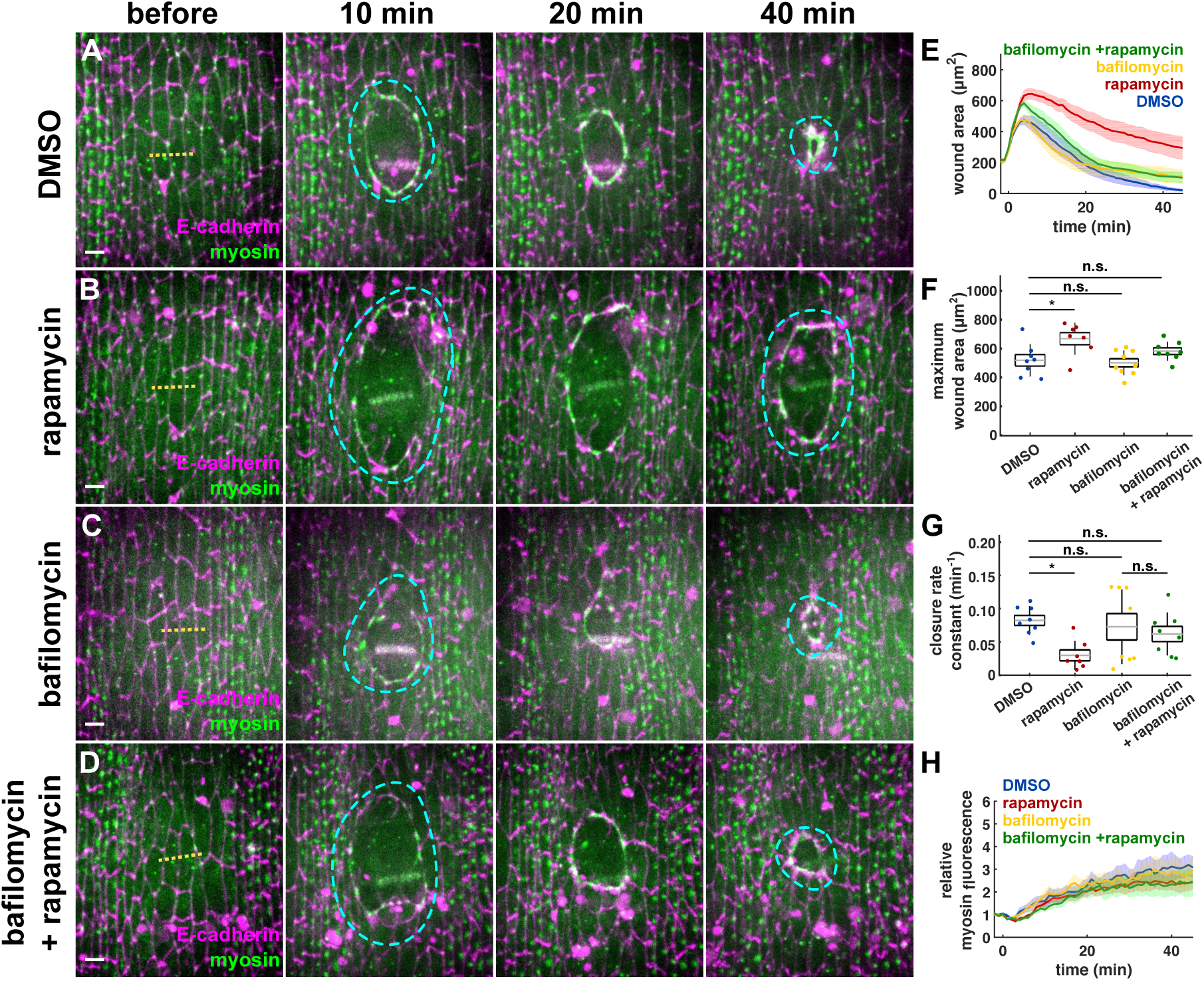
Acute autophagy inhibition when mTOR is disrupted rescues rapid wound healing. **(A-D)** Wound closure in embryos expressing myosin:GFP (Sqh, green) and E-cadherin:tdTomato (magenta), and injected with DMSO (A), rapamycin (B), bafilomycin (C), or both rapamycin and bafilomycin (D). Yellow dashed lines indicate wound sites, cyan dashed lines outline wounds. Time after wounding is shown. Anterior left, dorsal up. Bars, 5 μm. **(E-H)** Wound area over time (E), maximum wound area (F), wound closure rate constant (G), and relative mean myosin fluorescence at the wound edge (H) for embryos injected with DMSO (blue, *n* = 8 wounds), rapamycin (red, *n* = 7), bafilomycin (yellow, *n* = 7), or co-injected with rapamycin and bafilomycin (green, *n* = 8). (E, H) Error bars, s.e.m.. (F-G) Error bars, s.d.; boxes, s.e.m.; gray lines, mean. n.s., not significant; * *P* < 0.05.

